# Efficient production of functional proaerolysin from *E.coli*

**DOI:** 10.1101/2024.09.19.613822

**Authors:** Quynh Thi-Huong Pham, Ayako Tagawa, Narumi Iwata, Yusuke Miyanari

## Abstract

Proaerolysin is a bacterial toxin produced by *Aeromonas hydrophila* that specifically binds to GPI-anchored proteins on the plasma membrane, creating transmembrane pores that lead to cell death within a few hours. Leveraging this unique property, proaerolysin is widely used in diagnostic tests for paroxysmal nocturnal hemoglobinuria (PNH), a disease caused by somatic mutations in the *PIGA* gene, which is involved in the biosynthesis of GPI anchors. Additionally, proaerolysin serves as a counter-selection agent in genetic manipulations. Although bacterial expression and purification of proaerolysin have been previously reported, yields were low due to the absence of internal disulfide bonds, which are crucial for protein stability. Here, we demonstrate that using the Shuffle *E. coli* strain, which facilitates the formation of disulfide bonds in the cytoplasm, significantly improves the solubility and proper folding of proaerolysin. We achieved a high yield of proaerolysin, approximately 3 mg from a 50 ml bacterial culture, with a purity of over 99%. The functionality of recombinant proaerolysin was confirmed by testing in mouse embryonic stem cells (mESCs), demonstrating that this high-yield production method offers a reliable and cost-effective source of functional proaerolysin for a wide range of biotechnological applications.

## Introduction

Proaerolysin is a bacterial prototoxin produced by *Aeromonas hydrophila*, which selectively binds to glycosylphosphatidylinositol (GPI)-anchored proteins on the plasma membrane (Abrami et al., 1998). Upon binding, it perforates the membrane by forming transmembrane channels, causing ion imbalances and osmotic swelling, and ultimately leading to cell death within a few hours (Fig. 1a). Due to its membrane-disrupting mechanism, proaerolysin has been widely studied, serving not only as a model system for understanding pore-forming toxins (Abrami et al., 2000; Cirauqui et al., 2017; Etxaniz et al., 2018) but also for its potential applications in diagnostic and basic research (Brodsky et al., 2000; Cao et al., 2019). For example, fluorescently labeled proaerolysin (FLAER) has been employed as a diagnostic tool for detecting GPI-anchored proteins in paroxysmal nocturnal hemoglobinuria (PNH), a disease primarily caused by acquired somatic mutations in the *PIGA* gene (Manivannan et al., 2020). The mutations in *PIGA*, an essential enzyme for GPI anchor synthesis, cause the general deficiency of GPI-anchored proteins on blood cells (Ware et al., 1994), leading to intravascular hemolysis, blood clots, and bone marrow failure. The absence of the fluorescent signals of FLAER on blood cells allows for the detection of PNH through flow cytometry, making it a highly sensitive and accurate diagnostic method (Manivannan et al., 2020).

**Fig 1.**
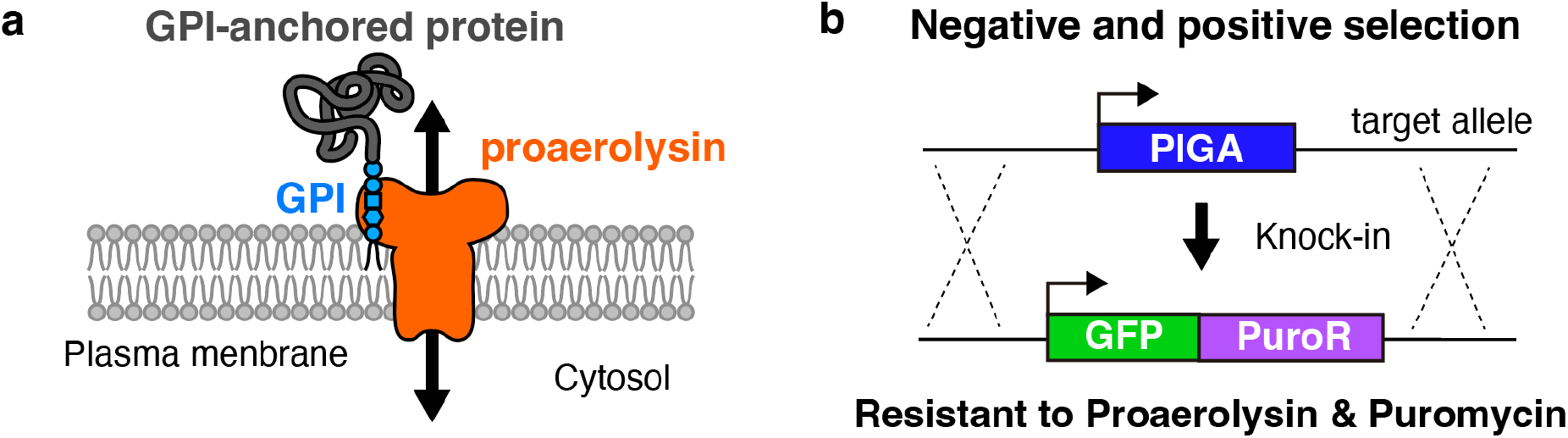
Proaerolysin and its application. **a**) Schematic representation of the molecular action of proaerolysin to GPI-anchored protein, leading to perforation of the plasma membrane. **b**) Example of positive and negative selection using *PIGA* and *PuroR*. The *PIGA* gene, a negative selection marker pre-inserted at the target locus, is replaced by the positive selection marker GFP-Puro through knock-in.

Leveraging the PIGA-dependent toxicity, proaerolysin has recently been utilized for negative selection in advanced genome editing techniques such as BIG-IN (Brosh et al., 2023, 2021; Li et al., 2021; Zhang et al., 2022). Pre-insertion of the *PIGA* gene at a target locus as a negative selection marker ensures efficient recombination using a targeting vector containing a positive selection marker, such as the Puromycin resistance gene (Fig. 1b). The combination of positive and negative selection enhances the accuracy of genome editing, making it particularly effective for large-genome rewriting (Brosh et al., 2023, 2021; Ordoñez et al., 2024; Zhang et al., 2023). Although the PIGA/proaerolysin system requires deletion of the endogenous *PIGA* gene on the X chromosome, it offers significant advantages over traditional negative selection markers, such as HSV-TK and HPRT. Notably, the PIGA/proaerolysin system, unlike others, does not exhibit the bystander effect, which unintentionally eliminates the neighboring cells (Brosh et al., 2021). Additionally, proaerolysin is highly efficient, killing cells within a few hours, while other systems require several days to achieve the same effect.

The efficient preparation of proaerolysin is a key step for several applications. Although bacterial expression of proaerolysin with BL21 E.coli strain was reported previously (Zhang et al., 2013), we found that the solubility of proaerolysin in BL21 was poor, possibly due to the lack of internal disulfide bonds. In this study, we used Shuffle strain to overcome the issue and achieved highly efficient and reliable purification of proaerolysin. We confirmed the recombinant proaerolysin is functional and applicable for negative selection for genome editing in mouse embryonic stem cells (mESCs).

## Results

### Preparation of soluble proaerolysin using a Shuffle strain

We constructed a bacterial expression plasmid, pET-proaerolysin-cH10, encoding proaerolysin with a His-tag fused to the C-terminus. The proaerolysin cDNA was optimized for *E. coli* codon usage and for reduced mRNA secondary structure near the start codon to enhance translation efficiency (Bhandari et al., 2021). As an initial trial, we transformed the plasmid into *E*.*coli* BL21(DE3) and induced protein expression with IPTG at 18°C overnight. We extracted proteins from cells using chemical lysis and sonication, and evaluated protein levels in both whole-cell extracts and soluble fractions using SDS-PAGE with Coomassie Brilliant Blue (CBB) staining. Although proaerolysin was highly expressed, only a portion of the protein was solubilized, with the majority remaining insoluble (Fig. 2a). Notably, native proaerolysin contains two internal disulfide bonds that strongly stabilize the protein (Lesieur et al., 1999). Since *E. coli* BL21(DE3) cannot form disulfide bonds in the cytoplasm, we hypothesized that the lack of these disulfide bonds in recombinant proaerolysin caused misfolding, contributing to its poor solubility. To address this issue, we tested the Shuffle strain, an engineered *E. coli* line that expresses disulfide bond isomerase DsbC in the cytoplasm, which promotes disulfide bond formation and proper protein folding (Lobstein et al., 2012). Remarkably, expression in the Shuffle strain significantly improved the solubility of proaerolysin compared to BL21(DE3) (Fig. 2a). Therefore, proaerolysin expressed in the Shuffle strain was selected for subsequent purification. We initially performed immobilized metal affinity chromatography (IMAC) to purify the His-tagged proaerolysin, achieving a purity of 84% (Fig. 2b, 2d). The IMAC eluate was subsequently purified using size exclusion chromatography. We observed a prominent main peak at 14 ml, accompanied by a small shoulder around 15 ml, both of which contained proaerolysin (Fig.2c, 2d). Calculation based on size markers showed that the main peak corresponds to 73.2 kDa, which is nearly double that of the shoulder, approximately 37.7 kDa. Given that proaerolysin is known to form dimers (Barry et al., 2001; Fivaz et al., 1999; Van Der Goot et al., 1993), it is likely that the main peak represents the dimer and the shoulder represents the monomer. However, there is a slight difference in the actual molecular weight of His-tagged proaerolysin, which is 53.7 kDa. Since the dimerization of proaerolysin enhances the protein stability, we selected fractions of the main peak as the final product. SDS-PAGE analysis showed a mobility shift of proaerolysin upon treatment of reducing reagent 2-mercaptoethanol, confirming the formation of intermolecular disulfide bonds in the purified proaerolysin (Fig. 2e). As a result, we obtained approximately 3 mg of proaerolysin, with a purity of 99.3%, from a 50 mL bacterial culture.

**Figure 2.**
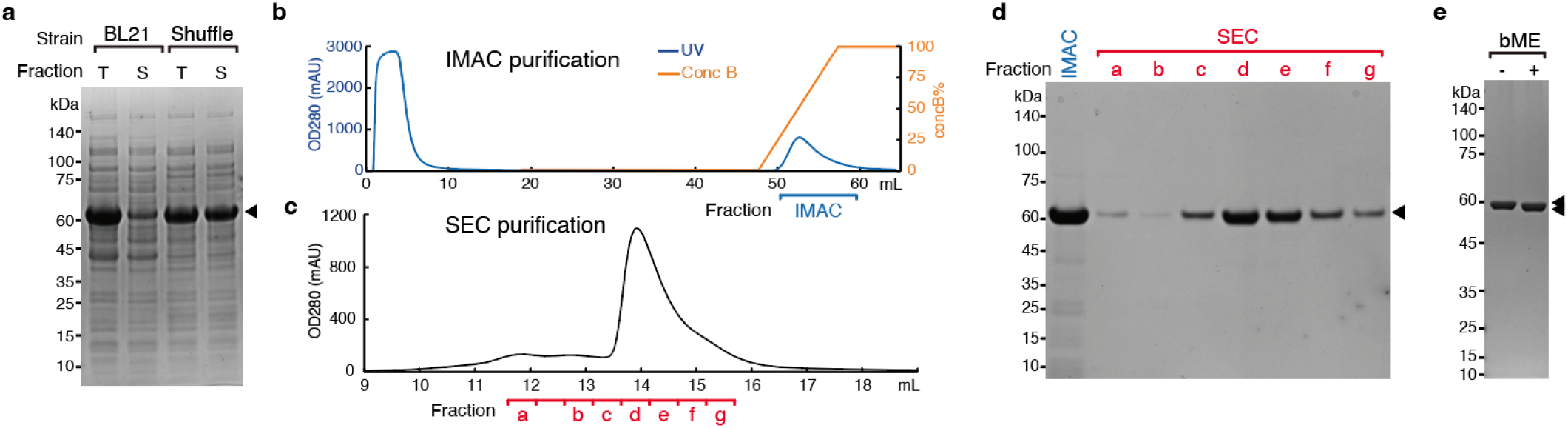
Purification of proaerolysin expressed in Shuffle. **a**) An image for CBB staining of an SDS-PAGE gel analyzing the total cell extracts (T) and soluble fractions (S) from BL21(DE3) and Shuffle strains expressing proaerolysin. The molecular weight marker sizes (in kDa) are indicated on the left. The band corresponding to proaerolysin is marked with an arrowhead. **b, c**) Profiles of IMAC (b) and SEC (c) purification are displayed. The elution fractions analyzed by SDS-PAGE are indicated at the bottom of each profile. **d**) A CBB-stained SDS-PAGE gel analyzing the elution fractions indicated in b and c. **e**) CBB staining of an SDS-PAGE gel analyzing the final purified proaerolysin with or without beta-mercaptoethanol treatment.

### The recombinant proaerolysin is functional

Proaerolysin specifically binds to GPI anchors on the plasma membrane, creating pores that result in rapid cell death (Diep et al., 1998). To evaluate the activity of our recombinant proaerolysin, we treated mESCs with varying concentrations of proaerolysin. The treatment induced drastic morphological changes within 2 hours, accompanied by cell detachment from the culture dishes (Fig. 3a). After 6 hours of treatment, cells were efficiently stained with trypan blue, a marker for plasma membrane permeability (Grankvist et al., 1979), in a concentration-dependent manner (Fig. 3a, b), indicating proaerolysin-induced membrane perforation and significant cell death. The concentration of proaerolysin required to efficiently induce cell death, particularly at 1 nM or higher, was comparable to that used in previous reports (Brosh et al., 2021; Zhang et al., 2022), supporting the functionality of proaerolysin. Upon treating the cells with 1 nM proaerolysin for 1 day, no recovery of mESCs was observed after an additional 3 days without proaerolysin, confirming its effectiveness as a selection agent. To further verify the integrity of proaerolysin, we generated *PIGA*-knockout mESCs using CRISPR/Cas9 with two guide RNAs targeting the 5’ and 3’ regions of the *PIGA* gene, resulting in the deletion of the entire *PIGA* gene locus (Fig. 3c). Since the loss of *PIGA* is expected to impair the GPI-anchoring process, thereby conferring resistance to proaerolysin (Diep et al., 1998). Genotyping PCR with a primer pair flanking the target region (17.8 kb in the wild type) produced a 900 bp amplicon in the *PIGA*-knockout, while the wild-type allele could not be amplified due to its large size, confirming the deletion of the *PIGA* gene in this cell line (Fig. 3c). We then tested the *PIGA*-knockout mESCs with up to 16 nM of proaerolysin for 24 hours and observed that the cells were well resistant to the toxin and continued to proliferate normally (Fig. 3d), in contrast to the severe cell death observed in parental mESCs. These results suggest that the cell death induced by the treatment occurred through its natural mechanism of action, suggesting that recombinant proaerolysin is functional.

**Figure 3.**
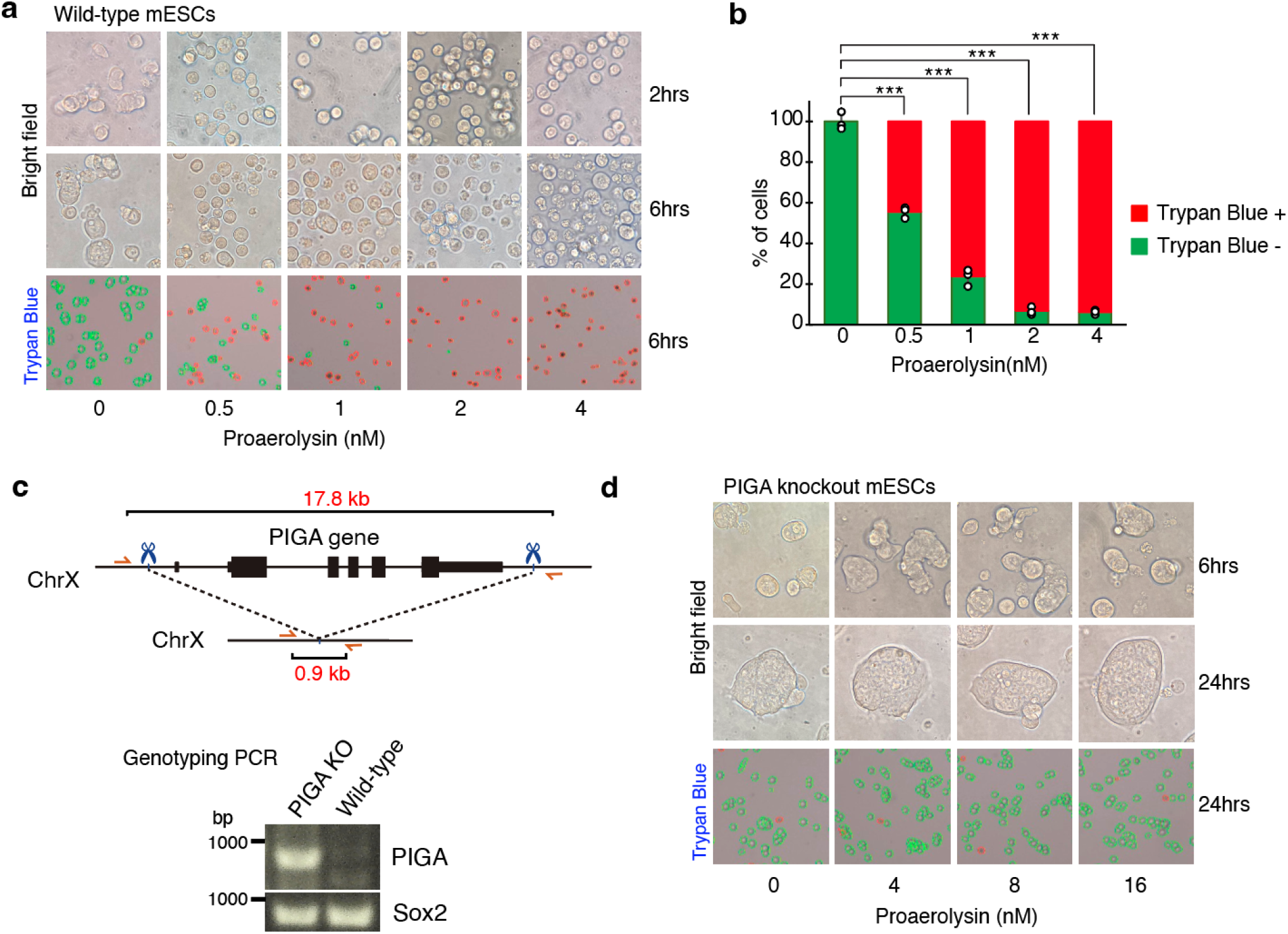
Functional validation of recombinant proaerolysin using mESCs. **a**) Wild-type mESCs were treated with indicated concentrations of proaerolysin(0, 0.5, 1, 2, and 4 nM). Images for bright-field microscopy (top and middle rows) and trypan blue staining (bottom row) are shown. The treatment durations (2 and 6 hours) are indicated on the right side of the corresponding rows. In the trypan blue staining, viable cells without staining are marked with green circles, while dead cells with staining are marked with red circles. **b**) The percentage of cells with or without trypan blue staining is shown. Individual data points are displayed as dots (n=3). Statistical significance was analyzed using a two-tailed t-test, with ^***^ representing p < 0.001. **c**) The top diagram shows the genomic locus of the mouse PIGA gene before and after CRISPR/Cas9-mediated knockout. The positions targeted by sgRNAs are marked with scissors, and the annealing sites for the primers used in genotyping PCR are indicated by orange arrows. The expected sizes of PCR products are shown (17.8 kb and 0.9 kb, respectively). The wild-type allele could not be amplified by genotyping PCR due to its large size. The bottom panels display agarose gel images of the genotyping PCR results. Sox2 is used as a control locus. **d**) PIGA knockout mESCs were treated with indicated concentrations of proaerolysin. Images for bright field and trypan blue staining are shown as **a**.

## Discussion

In this study, we efficiently expressed soluble proaerolysin using the SHuffle strain. The functionality of the purified protein was confirmed by assessing its perforating activity in wild-type and *PIGA*-knockout mESCs. While the BL21 strain was previously used to express recombinant proaerolysin (Zhang et al., 2013), its solubility was likely poor, possibly due to the absence of two internal disulfide bonds that stabilize the proaerolysin protein (Lesieur et al., 1999). Production of recombinant proteins with proper disulfide bonds, which are particularly important for protein stability and folding, is challenging. Although in vitro refolding of proteins with oxidizing reagents can promote the formation of disulfide bonds, it is often accompanied by protein aggregation due to misfolding or improper disulfide bond formation, resulting in low yield, and thus requires extensive optimization (Huang et al., 2012; Singh et al., 2015). Expression of proteins in the E. coli periplasm, where the Dsb protein family facilitates disulfide bond formation, offers an alternative approach. However, this approach typically results in lower protein yields compared to cytoplasmic production and requires the addition of an N-terminal signal peptide for periplasmic targeting (Huang et al., 2012). The Shuffle strain overcomes these limitations by enabling disulfide bond formation in its cytoplasm, as demonstrated by the mobility shift on SDS-PAGE under a reducing condition (Figure 2e). The Shuffle is a genetically modified *E*.*coli* strain lacking thioredoxin reductase (trxB) and glutathione reductase (gor) pathways, coupled with the cytoplasmic expression of DsbC (Bessette et al., 1999; Lobstein et al., 2012). Particularly, DsbC possesses dual activities: correcting improperly oxidized proteins and functioning as a chaperone to assist in protein folding (Chen et al., 1999). Thanks to these advantages, the Shuffle strain has been successfully used to express various disulfide-bonded proteins as soluble and functional proteins, such as Tissue Plasminogen Activator and nanobodies (Lobstein et al., 2012; Pleiner et al., 2015). In this study, we demonstrated the effective production of proaerolysin using the Shuffle strain, achieving a yield of 3 mg of highly purified protein with 99% purity from a 50 ml bacterial culture.

Negative selection markers are invaluable for genome rewriting through homologous recombination due to their capacity for efficient selection against large cell populations. Traditional systems like HSV-TK/ganciclovir and HPRT/6-thioguanine are widely used, partly due to the availability of commercial selection drugs. However, these two systems accompany bystander effects or spontaneous mutations that compromise counterselection accuracy (Brosh et al., 2021; Huo et al., 2001). The PIGA/proaerolysin system has emerged as a promising alternative, offering high selection efficiency without bystander effects (Brosh et al., 2021). Nonetheless, its applications require the pre-knockout of the endogenous X-linked *PIGA* gene and PIGA is essential for mouse development. This issue was addressed by developing a conditional system that allows precise control of *PIGA* expression through the introduction of a Cre-mediated STOP cassette into the intron of the *PIGA* gene (Zhang et al., 2022). This cassette blocks PIGA expression until Cre recombinase is applied, which removes the STOP cassette and allows PIGA to be re-expressed. Another challenge to the widespread use of this system is the limited commercial availability of proaerolysin, with the only widely available modified form being FLAER, fluorescent proaerolysin for diagnosing PNH. Therefore, our optimized production of recombinant proaerolysin overcomes this supply issue and broadens the scope for its use beyond genome editing, including applications such as nanopore sensing (Cao et al., 2019).

Proaerolysin is proteolytically cleaved to release a C-terminal peptide, converting it into its active form, aerolysin (van der Goot et al., 1994). Compared to the active form, proaerolysin is much less toxic to mice and humans (Bernheimer et al., 1975; Howard and Buckley, 1985), making it significantly safer for experimental handling. Nonetheless, it’s important to ensure lab workers are properly protected and to follow basic safety rules when working with proaerolysin, as it remains a toxic compound.

## Figures

## Materials and Methods

### Construction of plasmid DNA expressing His-tagged proaerolysin

The cDNA of the proaerolysin sequence (UniProt ID P09167) was synthesized by IDT, with *E. coli* codon usage and the 5’ mRNA region optimized using IDT Codon Optimization Tool (idtdna.com/CodonOpt) and TISIGNER (Bhandari et al., 2021), respectively. The DNA fragment was directly cloned into the pET28-cH10_ccdB vector by Gibson assembly using NEBuilder (NEB), resulting in the pET-proAER-H10 construct. The sequence of the insert was confirmed by Sanger sequencing (Eurofin Genomics). The pET-proAER-H10 is deposited to Addgene.

### Expression and purification of proaerolysin

pET-proaerolysin-cH10 was transformed into a chemically competent *E*.*coli* BL21(DE3) (Nippongene, 314-06533) or Shuffle T7 Express lysY (NEB, C3030). The cells were plated on LB agar supplemented with 50 µg/mL kanamycin and then cultured at 37 °C overnight until colonies were observed. A single colony was inoculated into 2mL of 2YTK (16 g of tryptone, 10 g of yeast extract, and 5 g of NaCl per 1L, containing 20 µg/ml of kanamycin) and incubated at 37°C overnight with shaking at 200 rpm. 1 ml of the pre-culture was inoculated into 50 ml of 2YTK and cultured at 37°C until A_600_ 0.6. The protein induction was then started by adding 0.1 mM isopropyl β-D-1-thiogalactopyranoside (IPTG). The cultures were grown further with shaking at 200 rpm at 18°C overnight and harvested by centrifugation at 4000×g for 15 min at 4°C. The cell pellet was stored at -20°C until purification. All purification steps were performed at 4°C. The *E*.*coli* cells were resuspended in 3 ml of IMAC A1 buffer (20 mM HEPES pH8.0, 500 mM NaCl) supplemented with 10 µg/ml DNase I, 1 µg/ml RNase A, 20 µg/ml Lysozyme, 0.2mM PMSF, and 1x CellLytic B Cell Lysis Reagent (Sigma, C8740) and then incubated at 37°C for 5 min. The cell lysate was sonicated using a BioRuptor (CosmoBio) at high power with 30 seconds ON and 30 seconds OFF cycles for 10 min. The BL21(DE3) lysate remained cloudy even after the sonication due to the abundant insoluble fraction, in contrast to the cleared lysate of Shuffle. The lysate was centrifuged at 15000 rpm for 10 min to remove the insoluble fractions. The cleared lysate was then filtered through a 0.45 µm syringe filter and applied to 1 mL of HisTrap Excel (Cytiva) for affinity purification using an AKTA-go system (Cytiva). The column was washed with 30 column volumes of IMAC A buffer containing 10 mM imidazole at a flow rate of 1.0 mL/min. His-tagged proaerolysin was eluted using a linear gradient up to 100% IMAC B buffer (IMAC A buffer containing 1 M imidazole) over 10 column volumes. The eluate was concentrated to 1000 µl by ultracentrifugation using a 10kDa VivaSpin Turbo 15 concentrator (Sartorius, VS15T02). After filtering the sample using a 0.22 µm spin filter (Protein Ark, GEN-MSF500), 500 µl of the sample was applied to a Superdex 200 increase 10/300 GL column (Cytiva) for size-exclusion chromatography. The mobile-phase buffer was TBSE (20 mM Tris-HCl pH 8.0, 150 mM NaCl, 1 mM EDTA) and the applied flow rate was 0.5 mL/min. The peak fractions containing purified proaerolysin were combined and its protein concentration was determined based on UV absorption at 280 nm with the extinction coefficient of 130290 cm-1 M-1 or 0.412 mg/mL (7.66 µM) for OD280 = 1. The SEC purification was repeated again for the remaining 500 µl of the IMAC eluate. The yield of purified proaerolysin was approximately 3 mg with 99% purity from a 50 mL of C3030 bacterial culture. The sample was sterilized by filtration using a 0.22 µm spin filter (Corning, 8160) and stored at -80°C. For practical use, the samples were diluted to 5 µM with OptiMEM (ThermoFisher), and 100 µL aliquots were stored at -80°C or 4°C for short-term use.

### SDS-PAGE

SDS-PAGE was performed as described previously (Schägger and Von Jagow, 1987). Briefly, SDS-PAGE was conducted using a Bis-Tris 4-10% gradient or 10% polyacrylamide gel with MES or MOPS running buffer. After running, the gel was stained with Coomassie Brilliant Blue R-250. The CBB-stained gel was scanned using Chemi-Doc (Bio-Rad). The quantity and purity of proaerolysin was calculated by using Image Lab (Bio-Rad). To assess the presence of potential disulfide bonds, proaerolysin samples were prepared in the LDS sample loading buffer with or without 2-mercaptoethanol and incubated at 95°C for 5 min before running.

### Cell culture

mESCs (E14) were cultured in DMEM containing 20% FBS, leukemia inhibitory factor, 3 mM CHIR99021 (a GSK3b inhibitor, Sigma), 1 mM PD0325901 (a MEK inhibitor, Sigma), 1 mM sodium pyruvate, penicillin/streptomycin, non-essential amino acids, and 0.1 mM 2-mercaptoethanol. The cells were grown at 37°C with 5% CO_2_.

### Preparation of sgRNA

A pair of small guide RNAs (sgRNAs) targeting the mouse *PIGA* genes (∼17 kb) was selected from a previous publication (Brosh et al., 2021), with the sequences 5’-GGCATGCTTTGTGGTCGTTC-3’ and 5’-CCCGCGGGCAGCCTATATAA-3’. Preparation of *in vitro* transcribed sgRNAs was performed as described before (Ishii et al., 2024). Briefly, template DNA for in vitro transcription (IVT) was prepared via two rounds of PCR. The first PCR was conducted using a primer pair (sgR-F1; GTTTGAGAGCTATGCTGGAAAC, sgR-R1: AAAAGCACCGACTCGGTGCC), a template DNA pU6-sgR1.2, and KOD FX Neo PCR Reagent (Toyobo, KFX-201). After DpnI treatment, the PCR product was purified with Monarch PCR DNA Cleanup Kit (NEB, T1030S) and diluted to 10 ng/uL. The second PCR was performed using a T7 promoter-gRNA-containing forward primer (PG.KO-sgRF1: TTCTAATACGACTCACTATAGGCATGCTTTGTGGTCGTTCGTTTGAGAGCTATGCTGGAAAC or PG.KO-sgRF2:TTCTAATACGACTCACTATAGGCCCGCGGGCAGCCTATATAAGTTTGAGAGCTATGCT GGAAAC) and the reverse primer sgR-R1. After the purification, the PCR products were diluted to 50 ng/uL and used as templates for subsequent IVT reactions. The sgRNA was transcribed *in vitro* overnight at 37°C using the CUGA in vitro Transcription Kit (Nippon Gene, 307-13531) following the manufacturer’s instructions. Post-transcription, the RNA was treated with DNase I at 37°C for 30 minutes to remove the template DNA and then purified via ammonium acetate precipitation. The transcribed RNA was confirmed by agarose gel electrophoresis, quantified using the Qubit RNA BR Assay Kits (ThermoFisher, Q10210), and adjusted to 1 µg/µL.

### Knockout of PIGA gene in mouse ES cells

Knockout of the *PIGA* gene in mES cells was performed by electroporation of the CRISPR-Cas9/sgRNA ribonucleoprotein (RNP) complex, as previously described (Brosh et al., 2021). Briefly, the Cas9 RNP complex was prepared by mixing 100 pmol of a pair of gRNAs with 50 pmol of Cas9-NLS (Nippon, 316-08651) and 40 mg/mL poly(glutamic acid) (Sigma, P4761-100MG) (Nguyen et al., 2020). After incubating at 37°C for 15-30 minutes to allow RNP complex formation, the Cas9/RNP complex was combined with 100 µL of OptiMEM I medium (Thermo, 31985-062) containing 1 × 10^6^ cells, and then electroporated using the CUY21EDIT II (BEX) electroporator under the following conditions: 125V, 1 msec ON, 50 msec OFF, with 5 pulses. Immediately after electroporation, 400 µL of culture medium was added to the cuvette, and the cell suspension was transferred to a 6-well plate with an additional 2 mL of culture medium. 7 days post-electroporation, cells were selected by treating with 2-4 nM of proaerolysin for 2 days.

Surviving cells were then subjected to genotyping PCR to verify the deletion of the entire *PIGA* gene locus. Briefly, genomic DNA was extracted using a Genome Lysis Buffer (20 mM Tris pH 8.0, 10 mM EDTA, 100 mM NaCl, 0.5% SDS, and 5 µg/ml Proteinase K), and then purified by isopropanol precipitation. Genotyping PCRs were performed using KOD One PCR Master Mix -Blue- (Toyobo, KMM-201), 300-400 ng of genomic DNA, and the following primer pairs: mPIGA.genoF3 (TGTGAGGTGCCTCCCACTCT) and mPIGA.genoR3 (TAGCAGAGCCCATGAATCCG) for *PIGA* gene locus, and mSox2.F (GGCGTGAACCAGCGCATGGAC) and mSox2.R (CAATGGACATTTGATTGCCATGT) for *Sox2* gene locus. PCR conditions included an initial denaturation at 95°C for 3 minutes, followed by 7 touch-down cycles (98°C for 5 seconds, 67°C for 30 seconds with decreasing 1°C per cycle, 68°C for 30 seconds), then 30 cycles of 98°C for 5 seconds, 61°C for 30 seconds, 68°C for 30 seconds. Control reactions amplifying the *Sox2* locus were included to verify the quality of genomic DNA. PCR products were analyzed using electrophoresis and confirmed by Sanger sequencing.

### Treatment of mESCs with proaerolysin

Wild-type and *PIGA* knockout mES cells were seeded at a density of 2 × 10^5^ cells per well into the 24-well plate coated with gelatin and cultured in 500 µl of culture medium containing the indicated concentration of proaerolysin. The effect of proaerolysin on cell viability was briefly assessed by observing cell morphology under microscopy. To evaluate the sensitivity of cells against proaerolysin, all cells were treated with trypsin 6 hours post-treatment, stained with Trypan Blue (Gibco, 15250-061), and quantified using Countess II Automated Cell Counter (Invitrogen).

## Acknowledgments

We appreciate all members of Miyanari lab for their kind discussion. Y.M. is supported by JSPS KAKENHI (21H04765, 23K21749, 22H04688), World Premier International Research Center Initiative (WPI, MEXT), Astellas Foundation, Takeda Science Foundation, The Mitsubishi Foundation, The Hokuriku Cancer Foundation, and the Naito Foundation. Q.T.H.P. is supported by JST SPRING, Grant Number JPMJSP2315.

## Author contributions

Q.T.H.P. and Y.M. designed experiments and wrote the paper. Q.T.H.P. performed all the experiments using mESCs. A.T. conducted protein purifications. N.I. supported constructions of plasmid DNAs and induction of protein expressions.

